# Acquisition of cell migration defines NK cell differentiation from hematopoietic stem cell precursors

**DOI:** 10.1101/142380

**Authors:** Barclay J. Lee, Emily M. Mace

## Abstract

Human natural killer (NK) cells are generated from CD34^+^ precursors and can be differentiated *in vitro* by co-culture with developmentally supportive stromal cells. Despite the requirement for stromal cell contact in this process, the nature of these contacts has been poorly defined. We have previously identified a requirement for NK cell signaling receptors associated with terminal maturation in NK cell migration. However, the relationship between NK cell migration and differentiation is still unclear. Here, we perform continuous long-term imaging and tracking of NK cell progenitors undergoing *in vitro* differentiation. We demonstrate that NK cell precursors can be tracked over long time periods on the order of weeks by utilizing phase-contrast microscopy, and show that these cells acquire increasing motility as they mature. Additionally, we observe that NK cells display a more heterogeneous range of migratory behaviors at later stages of development, with the acquisition of complex modes of migration that are associated with terminal maturation. Together these data demonstrate previously unknown migratory behaviors of innate lymphocytes undergoing lineage differentiation revealed by long-term imaging and analysis workflows.

## Introduction

Human natural killer (NK) cells are derived from CD34^+^ hematopoietic stem cell precursors that originate in the bone marrow and undergo terminal maturation in secondary lymphoid tissue (Freud *et al.*, 2005; Eissens *et al.*, 2012). Their differentiation has been stratified to 5 stages, with each stage marked by differential expression of surface markers, production of cytokines, and ability to perform cytotoxic functions (Freud *et al.*, 2005; Freud *et al.*, 2006). While originally these stages were identified in secondary lymphoid and tonsillar tissue, it is now appreciated that NK cells can likely undergo maturation in a variety of microenvironments within the body (Eissens *et al.*, 2012; Yu *et al.*, 2013). This has led to a model in which early NK cell precursors seed peripheral tissue and give rise to mature NK cells prior to entering circulation. Within peripheral blood, NK cells are found within the distinct functional and phenotypic subsets termed CD56^bright^ or CD56^dim^. These subsets are thought to represent discrete stages of the model described above, with multiple lines of evidence suggesting that the CD56^bright^ (Stage 4) subset is a direct precursor of terminally differentiated CD56^dim^ (Stage 5) NK cells (Chan *et al.*, 2007; Huntington *et al.*, 2009; Eissens *et al.*, 2012). However, given the seeming plasticity in environments that can support NK cell development, as well as the poor correlation between human and murine NK cell phenotypes, human NK cell differentiation is a poorly understood process. Human NK cell development can be studied with an *in vitro* model, which is also of interest for the generation of clinical grade NK cells for immune therapy (Dezell *et al.*, 2012; Rezvani and Rouce, 2015). The most efficient method for *in vitro* differentiation of human NK cells from CD34^+^ progenitor cells is through co-culture with an irradiated feeder layer of stromal cells, specifically the EL08.1D2 cell line (Grzywacz *et al.*, 2006; Dezell *et al.*, 2012). *In vitro* derived NK cells have cytotoxic function and exhibit many of the same surface markers observed on mature NK cells isolated derived from patient blood (Miller and McCullar, 2001; Sivori *et al.*; Grzywacz *et al.*, 2006; Eissens *et al.*, 2012). NK cells can also be derived from progenitor cells in the presence of cytokines alone, but allowing contact with developmentally supportive stromal cells significantly increases cell proliferation as well as maturation (Miller and McCullar, 2001; Grzywacz *et al.*, 2006).

Despite the requirement for direct stromal cell contact in this system, the intercellular interactions between the CD34^+^ progenitors and the EL08.1D2 cells driving this developmental pathway are as of yet unclear and deserve further study. Incubation of CD56^bright^ or CD56^dim^ NK cells on EL08.1D2 stroma leads to significant non-directed migration on stromal cells. Furthermore, isolation of NK cell developmental intermediates (NKDI) from tonsillar tissue or *in vitro* differentiation identifies distinct migratory behavior based upon developmental stage (Mace *et al.*, 2016). Progression through differentiation is associated with increased velocity of migrating cells and decreased time spent in arrest. These migratory properties are conserved in both NKDI isolated from tonsillar tissue and NK cells generated *in vitro* from CD34^+^ precursors. (Mace *et al.*, 2016).

With current microscopy technology it is now feasible to perform long-term time-lapse imaging of cells with sufficient time resolution to adequately track cell movements. Many modern microscope systems allow for temperature and environmental regulation to ensure cell stability over a long period of time. Thus, a growing challenge in the field is how to best acquire and process large amounts of migration data in the most efficient and accurate manner. Typically, tracking is performed on cells labelled with a fluorescent dye, but this method is unsuited for long-term imaging, where phototoxicity and dye loss become problems (Coutu and Schroeder, 2013). Transmitted light microscopy avoids this issue, but makes tracking cells more difficult due to the lower signal-to-noise ratio in these images. In this case, tracking is often done manually, which is extremely time-consuming and allows for user bias. Reliable automated methods for tracking transmitted light images are thus highly desirable but currently lacking, although improvements have been made in recent years (Hilsenbeck *et al.*, 2016; Xing and Yang, 2016).

In this study, we show, for the first time, the continuous development of human NK cells from CD34^+^ hematopoietic stem cells (HSC) and demonstrate that the acquisition of motility is progressively acquired through NK cell maturation. We further quantify the heterogeneity in the developing NK cell migration phenotype by classifying single-cell tracks in terms of velocity and mode of migration (directed, constrained, or random). Together, this defines the acquisition of human NK cell migratory capacity throughout development using high temporal resolution and continuous length of imaging.

## Materials and Methods

### Cell culture

EL08.1D2 cells stromal cells (a gift from Dr. J. Miller, University of Minnesota) were maintained on gelatinized culture flasks at 32° C in 40.5% a-MEM (Life Technologies), 50% Myelocult M5300 (STEMCELL Technologies), 7.5% heat-inactivated fetal calf serum (Atlanta Biologicals) with β-mercaptoethanol (10^−5^ M), Glutamax (Life Technologies, 2 mM), penicillin/streptomycin (Life Technologies, 100 U ml^-1^), and hydrocortisone (Sigma, 10^−6^ M). Culture media was supplemented with 20% conditioned supernatant from previous EL08.1D2 cultures.

For *in vitro* CD34^+^ differentiation, 96-well plates were treated with 0.1% gelatin in ultrapure water to promote cell adherence. Gelatinized 96-well plates were pre-coated with a confluent layer of EL08.1D2 cells at a density of 5-10 x 10^3^ cells per well and then mitotically inactivated by irradiation at 300 rad. Purified CD34^+^ hematopoietic stem cells were cultured at a density of 2-20 x 10^3^ cells per well on these EL08.1D2 coated plates in Ham F12 media plus DMEM (1:2) with 20% human AB^-^ serum, ethanolamine (50 µM), ascorbic acid (20 mg l^-1^), sodium selenite (5 µg l^-1^), β-mercaptoethanol (24 µM) and penicillin/streptomycin (100 U ml^-1^) in the presence of IL-15 (5 ng ml^-1^), IL-3 (5 ng ml^-1^), IL-7 (20 ng ml^-1^), Stem Cell Factor (20 ng ml^-1^), and Flt3L (10 ng ml^-1^) (all cytokines from Peprotech). Half-media changes were performed every 7 days, excluding IL-3 after the first week.

NK92 cells (ATCC) were maintained in 90% Myelocult H5100 (STEMCELL Technologies) (Atlanta Biologicals) with IL-2 (200 U ml^-1^). YTS cells (a gift from Dr. J. Strominger, Harvard University) were maintained in 85% RPMI 1640 (Life Technologies), 10% fetal calf serum (Atlanta Biologicals), HEPES (Life Technologies, 10 µM), penicillin/streptomycin (Life Technologies, 100 U ml ^-1^), MEM Non-Essential Amino Acids Solution (Thermo Fisher, 1 mM), sodium pyruvate (1 mM), and L-glutamine (2 mM). All cell lines were authenticated by flow cytometry and confirmed monthly to be mycoplasma free.

### CD34^+^ precursor isolation and flow cytometry

T and B cell lineage depletion was performed using NK cell RosetteSep (STEMCELL Technologies) and Ficoll-Paque density gradient centrifugation from routine red cell exchange apheresis performed at Texas Children’s Hospital. Following pre-incubation with RosetteSep, apheresis product was layered on Ficoll-Paque for density centrifugation at 2,000 r.p.m. for 20 min (no brake). Cells were harvested from the interface and washed with PBS by centrifugation at 1,500 r.p.m. for 5 min then resuspended in fetal calf serum for cell sorting. All samples were obtained under guidance and approval of the Institutional Review Board of Baylor College of Medicine in accordance with the Declaration of Helsinki. T- and B- cell depleted cultures were incubated with antibodies for CD34 (clone 561, PE conjugate, BioLegend, 1:100) prior to sorting. FACS sorting was performed using a BD Aria II cytometer with an 85 µm nozzle at 45 p.s.i. Purity after sorting was >90%. Primary NK cells for short-term imaging were isolated with NK cell RosetteSep.

For FACS analysis of CD34^+^ intermediates, a 6-colour flow cytometry panel was performed on a BD Fortessa using antibodies for CD56 (Clone HCD56, Brilliant Violet 605, BioLegend, 1:200), CD3 (Clone UCHT1, Brilliant Violet 711, BioLegend, 1:200), CD16 (Clone 3G8, PE-CF594 conjugate, BD, 1:300), CD94 (Clone DX22, APC conjugate, BioLegend, 1:100), CD117 (Clone 104D2, PE/Cy7 conjugate, BioLegend, 1:10), and CD34 (clone 561, PE conjugate, BioLegend, 1:100). Flow cytometry data analysis was performed with FlowJo X (TreeStar Inc.).

### Live-cell imaging and tracking

Cells were imaged in 96-well ImageLock plates (Essen Bioscience) on the IncuCyte ZOOM Live-Cell Analysis System (Essen Bioscience) at 37° C every 2 minutes in the phase-contrast mode (10X objective). Images were acquired continuously for a 28-day period. NK cell lines and primary NK cells were labeled using CellTracker Violet (Thermo Fisher) at a working concentration of 15 µM and washed with complete media by centrifugation at 1,500 r.p.m. for 5 minutes. Fluorescent images were imported into Imaris (Bitplane) and tracked using the Spot tracking method with the ‘Autoregressive Motion Expert’ setting and removing tracks below an automatic threshold on ‘Track Duration’.

For tracking of NK developmental intermediates, cells were seeded at 2 x 10^3^ cells per well on a 96-well ImageLock plate with confluent irradiated EL08.1D2 cells, then imaged at 2 minute intervals for a 24-hour period. Phase-contrast images were segmented using ilastik software by using the ‘2-stage Autocontext’ method for pixel classification to distinguish NKDI from the background (Tu and Bai, 2010; Sommer *et al.*, 2011). An array of cell positions over time was generated from exported binary segmentation images and imported into Imaris (Bitplane) for tracking analysis. The image set was divided into 24 hour periods for tracking. Tracking was done using the Spot tracking method on Imaris with the ‘Autoregressive Motion’ setting and removing tracks below an automatic threshold on ‘Track Duration’. Individual tracks were then verified by eye and corrected manually where necessary by using Imaris to fix broken or overlapping tracks.

### Analysis

Track statistics were calculated using Imaris and exported as comma-separated value files which were graphed using GraphPad Prism. Mean speed was defined as the per-track average of all instantaneous speeds calculated at each frame. The straightness parameter was calculated by dividing the displacement by the total path length for each track, so that straightness values close to 1.0 represent highly directed tracks. Arrest coefficient was defined as the percentage of time that the cell stays in arrest based on a threshold on instantaneous speed of 2 µm/min, or approximately one cell body length per image interval. The standard deviation in velocity for each track was calculated as the standard deviation in instantaneous velocities observed for a given track. Rose plots were generated by selecting 30 representative tracks per time series and plotting them in Imaris.

For mean squared displacement (MSD) analysis and classification of migration modes, we developed custom MATLAB scripts based on a previously described method for transient migration behavior analysis (Khorshidi *et al.*, 2011). Calculation of MSD was done using @msdanalyzer, a publicly available MATLAB class for MSD analysis of particle trajectories (Tarantino *et al.*, 2014). To implement the method for transient migratory period analysis, each track was analyzed using a sliding window approach and calculating the MSD corresponding to each window. The MSD data was fit to curve to estimate the degree of curvature and fit to a line to estimate the diffusion coefficient of the cell. Track segments were then classified as directed, constrained, or random migration depending on thresholds on these values which were set to be the same as those previously described (Khorshidi *et al.*, 2011). The threshold for diffusion coefficient was set at 4.2 µm^2^ min^-1^, calculated based on the typical diameter of an NK cell, with all track segments having a smaller diffusion coefficient classified as constrained migration. On the other hand, all segments with an MSD curvature a above 1.5 were defined as directed migration. All remaining segments after these thresholds were applied were classified as diffusive migration.

All MATLAB code associated with this paper along with full documentation is available online on the project repository at https://github.com/barclaylee/Incucyte-analysis-toolbox.

### Statistics

Statistical analysis was calculated using Prism 6.0 (GraphPad). Ordinary one-way ANOVA was used to compare track statistics. Outliers were removed from exported track statistics using the ROUT method in Prism with Q = 1%. For all tests p < 0.05 was considered significant.

### Sample sizes

Given the varying number of tracks that met quality thresholds, the sample numbers were not equal for all samples tested. Specifically, the *n* for each representative experiment shown in the figures are: Fig. 1B: 75 (NK92), 205 (YTS), 250 (eNK); **Figs. 2B, 3A, 4A:** 932 (0D), 803 (7D), 134 (14D), 148 (21D); Fig. 2C, 3B: 607 (0D), 1052 (1D), 699 (2D), 419 (3D), 350 (4D), 372 (5D), 421 (6D), 376 (7D), 725 (8D), 903 (9D), 1022 (10D), 871 (11D), 712 (12D), 291 (13D); Fig. 4C: 932; Fig. 4D: 148.

**Figure 1.**
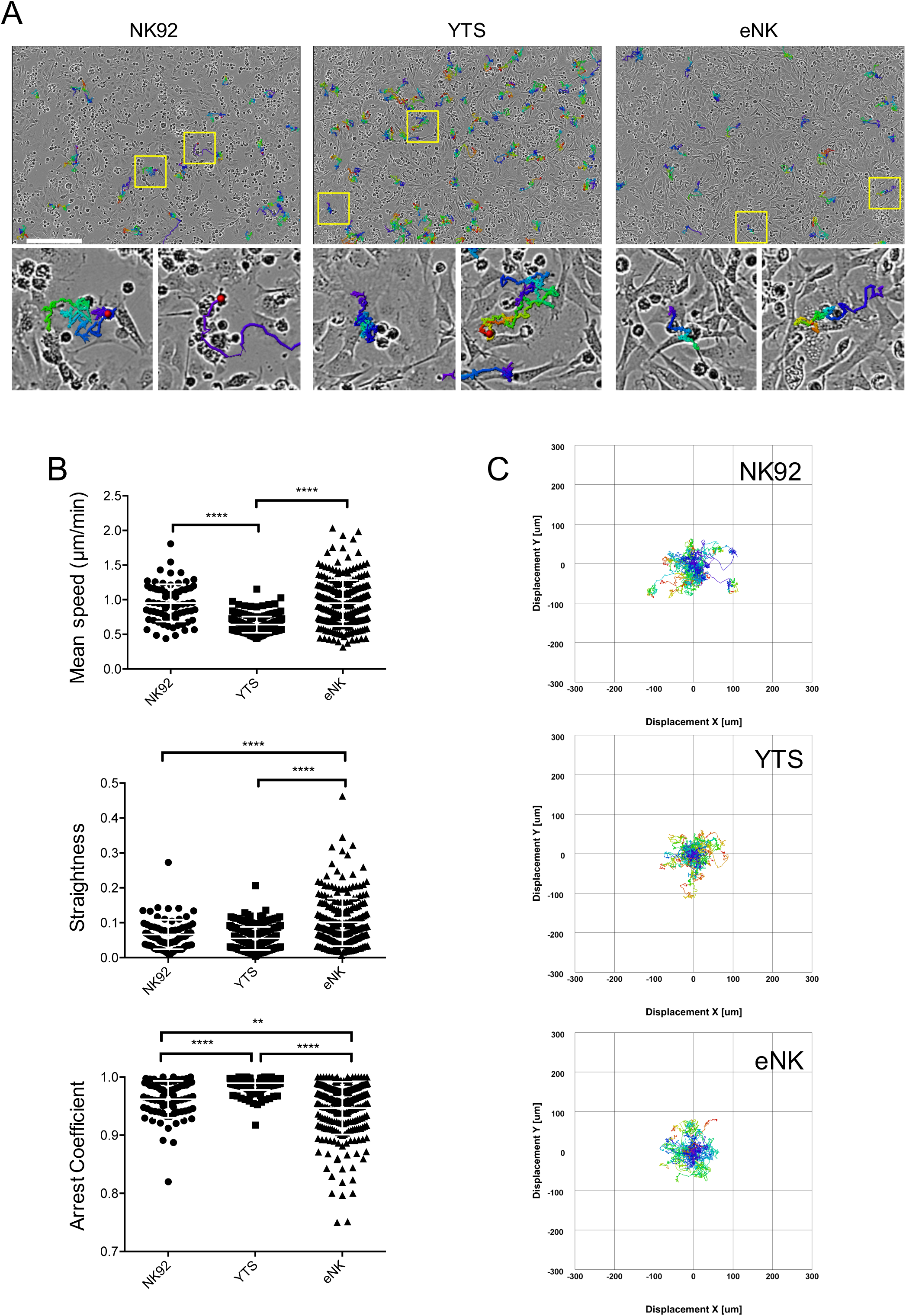
Continuous live cell imaging of human NK cells on stromal cells. YTS or NK92 NK cell lines or human NK cells (eNK) were labeled with CellTracker Violet then co-cultured with an irradiated EL08.1D2 monolayer for 24 hours in a 96-well plate. Images were acquired continuously every 2 minutes. A) Representative phase-contrast images of each cell type with randomly selected sample tracks overlaid. Insets show zoomed-in views of single-cell tracks. Scale bar=300 μm. B) Mean track speed (top), straightness (center), and arrest coefficient (bottom) of NK cells were calculated. Error bars indicate s.d. Means with significant differences were determined by ordinary one-way ANOVAwith Tukey’s multiple comparison test **p< 0.01, ****p< 0.0001). Data are representative of three independent experiments. n=75 (NK92), 205 (YTS), 250 (eNK). C) Rose plots of representative tracks. n=30 per graph.

### Results and Discussion

#### Human NK cell types exhibit distinct migratory behavior on stromal cells

NK cell lines and *ex vivo* NK cells undergo migration on EL08.1D2 stromal cells when assayed over relatively short periods of imaging (Mace *et al.*, 2016). To compare their migratory behavior and validate the sensitivity of our long-term imaging system, we labeled NK92 and YTS cell lines, and *ex vivo* NK cells (eNK) with CellTracker Violet dye, and co-cultured them with EL08.1D2 cells for 24 hours. Cells were continuously imaged at 2 minute intervals, and following acquisition cells were tracked in Imaris following segmentation based on fluorescence (Figure 1A). eNK and NK cell lines had distinct migratory properties classified by mean velocity, straightness, and arrest coefficient. eNK cells had more directed motion overall, characterized by significantly higher track straightness and lower arrest coefficient (frequency of time of imaging found in arrest) than both cell lines (Figure 1B, C). There were significant differences in motility between the two NK cell lines, with NK92 cells having higher overall speeds than YTS cells (0.953±0.27 µm/min vs. 0.6406±0.12 µm/min), although straightness was not significantly different (Figure 1B, C). This is in agreement with previous reports of NK92 and YT cells (from which the YTS line is derived) showing differential migration within Matrigel (Edsparr *et al.*, 2009). Collectively, these data demonstrate that the previously described migration of human eNK cells on stroma is recapitulated by NK cell lines, particularly with greatest fidelity by the NK92 cell line.

#### NK cell precursor motility and phenotype changes throughout maturation

As eNK cells show significant motility on stromal cells, and acquisition of migratory behavior is associated with NK cell development *in vitro* and *in vivo* (Mace *et al.*, 2016), we sought to define the migratory behavior of NK cell precursors throughout development. Specifically, we aimed to measure cell migratory properties with minimally invasive imaging in a large field of view in order to not bias analyses with cells that may leave the imaging field. Using our long-term imaging system described above, we generated human NK cells *in vitro* from purified CD34^+^ hematopoietic stem cells co-cultured on a developmentally supportive monolayer of EL08.1D2 stromal cells (Grzywacz *et al.*, 2006; Mace *et al.*, 2016). Cells were imaged continuously every 2 minutes throughout the 28-day period required for maturation into NK cells. Individual cells were segmented and tracked for parameters including track speed, length and displacement to measure how their migratory behaviors evolved throughout differentiation (Figure 2A). As expected, NK cell progenitors at the 0-day time point had significantly slower speeds than enriched NK cells (Figure 2B, 1B). All tracking was done on phase-contrast images because we found that fluorescent cellular dyes did not persist throughout the entire four-week period. Using this method enabled the tracking of single cells for extended periods of imaging (Supplemental Movie 1) and visual inspection confirmed that average cell track length increased significantly throughout the time of differentiation (Figure 2B, Supplemental Movies 2, 3).

**Figure 2.**
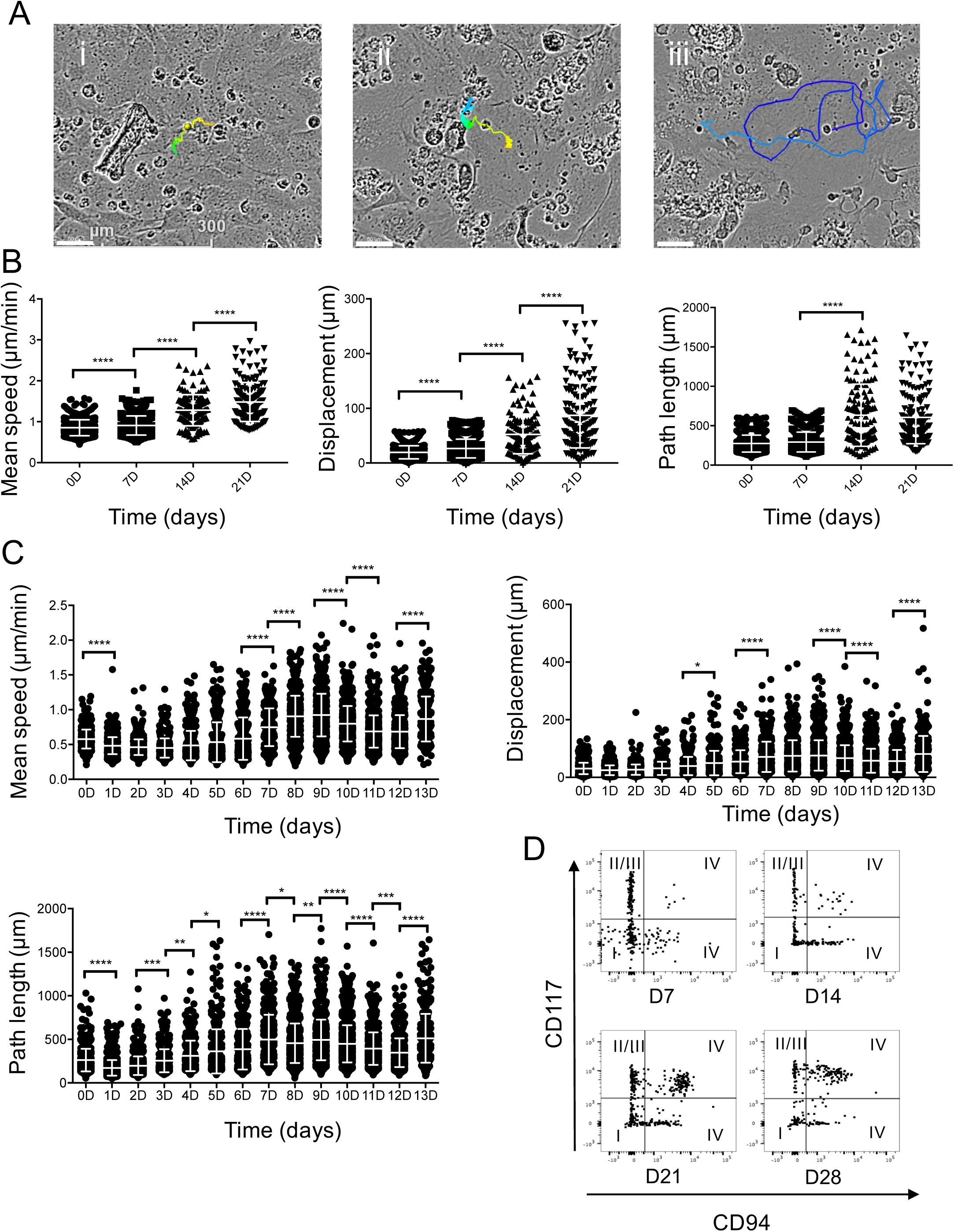
Acquisition of intrinsic NK cell migration with differentiation. CD34^+^ HSCs were seeded on a monolayer of EL08.1D2 cells. Cells were imaged continuously in phase-contrast mode for 28 days. FACS analysis was performed weekly to monitor expression of developmental markers. Representative images of NK cell intermediates with randomly selected tracks 0 (i), 7 (ii), and 21 (iii) days after start of experiment. Scale bar=50 mm. B) Mean speed, displacement, and path length of NK cell developmental intermediates were measured at 7-day intervals as indicated. Error bars indicate s.d. ****p< 0.0001 by ordinary one-way ANOVA with Tukey’s multiple comparisons test. n=932 (0D), 803 (7D), 134 (14D), 148 (21D). C) Mean speed, displacement, and path length of cells from continuous tracking from the first 14 days are shown as 24 hour segments. Error bars indicate s.d. Means with significant differences as analyzed by ordinary one-way ANOVA with Tukey’s multiple comparison test are shown (*p< 0.05, **p< 0.01, ***p< 0.001, ****p< 0.0001). Sample size for each individual time point are listed in Materials and Methods. D) FACS analysis of NK cell maturation markers. Predicted NK cell developmental stage based on phenotype as described in the text is shown in roman numerals. All data shown are representative of 3 independent experiments.

To quantify the migratory behavior of developing NK cells early in differentiation, tracks were extracted from continuous 24 hour periods at days 0, 7, 14 and 21. Mean velocity and displacement increased significantly at each progressive time point (Figure 2B). Initially, CD34^+^ cells had minimal velocity (0.58±0.14 µm/min), track length (255.58±131.07 µm), and displacement (31.91±21.54 µm). However, at day 21, mean velocity was 2.19±0.84 µm/min, consistent with previously reported migration speeds of mature NK cells (Khorshidi *et al.*, 2011; Mace *et al.*, 2016). Path length increased over time as well but was only significant between the 7-day and 14-day time points (Figure 2B). Given the significant differences between days 0 and 14, we performed an additional experimental replicate where cells were tracked for consecutive 24-hour movies over the first 14 days of imaging. Significant variations in mean speed, displacement, and path length were observed between daily time points, which cumulatively accounted for the overall trends previously described at the weekly time points (Figure 2C). Based on these results, we concluded that the acquisition of track speed and length in developing NK cells occurs progressively throughout the initial stages of differentiation. A similar increase in velocity has been similarly described in developing human T cells, as relatively immature (double-negative and double-positive) thymocytes have substantially slower track speeds than their more mature single-positive counterparts (Ehrlich *et al.*, 2009; Halkias *et al.*, 2013). On average, observed progenitor cell velocities at all stages were similar to typical velocities seen for immortalized NK cell lines as well as those seen in other studies looking at lymphocyte motility, both *in vitro* (Khorshidi *et al.*, 2011; Zhou *et al.*, 2017) and *in vivo* (Miller *et al.*, 2002; Ehrlich *et al.*, 2009).

Progression of NK cell development was additionally monitored by flow cytometry analysis of NK cell developmental markers at days 7, 14, 21, and 28. We analyzed cells for expression of CD117 and CD94, which, when considered with CD34, identify developmental stages 1-4, and CD16, a marker of NK cell terminal maturation (Freud *et al.*, 2006; Eissens *et al.*, 2012). NK cells undergo differentiation through four linear stages: stage 1 cells are CD34^+^CD117^-^CD94^-^, stage 2 cells are CD34^+^CD117^+^CD94^-^, stage 3 cells are CD34^-^CD117^+^CD94^-^, and stage 4 cells are CD34^-^CD117^+/-^CD94^+^. Increasing frequencies of cells underwent differentiation to stages 3 and 4 (35% at Day 28), which was accompanied by a decreasing frequency of immature cells at later time points (Figure 2D).

#### NK cell developmental intermediates in later stages of development exhibit more directed migration

Given the increasing speed and track length associated with migration of developing NK cells, we sought to further quantify the degree of directed migration in NK cell tracks by calculating the straightness and arrest coefficients. This was performed first on tracks extracted from 7-day intervals (Figure 3A). An increase in straightness and decrease in arrest coefficient over time was observed for the weekly time points (Figure 3A). While CD34^+^ HSC (0D) had a track straightness of 0.074±0.05, at the 21D time point the mean straightness index was 0.149±0.20. Similarly, arrest coefficient decreased from 0.993±0.01 to 0.884±0.1 between the 0 and 21-day time points. This was in agreement with the increase in mean speed observed for the later time points. As demonstrated for the significant changes in track velocity between days 14 and 21, straightness and arrest coefficient similarly had the greatest changes in this time interval (Figure 3A). Although significant differences from day to day were detected, the overall change in these statistics is relatively small compared to those observed between days 0 and 21 (Figure 3B).

**Figure 3.**
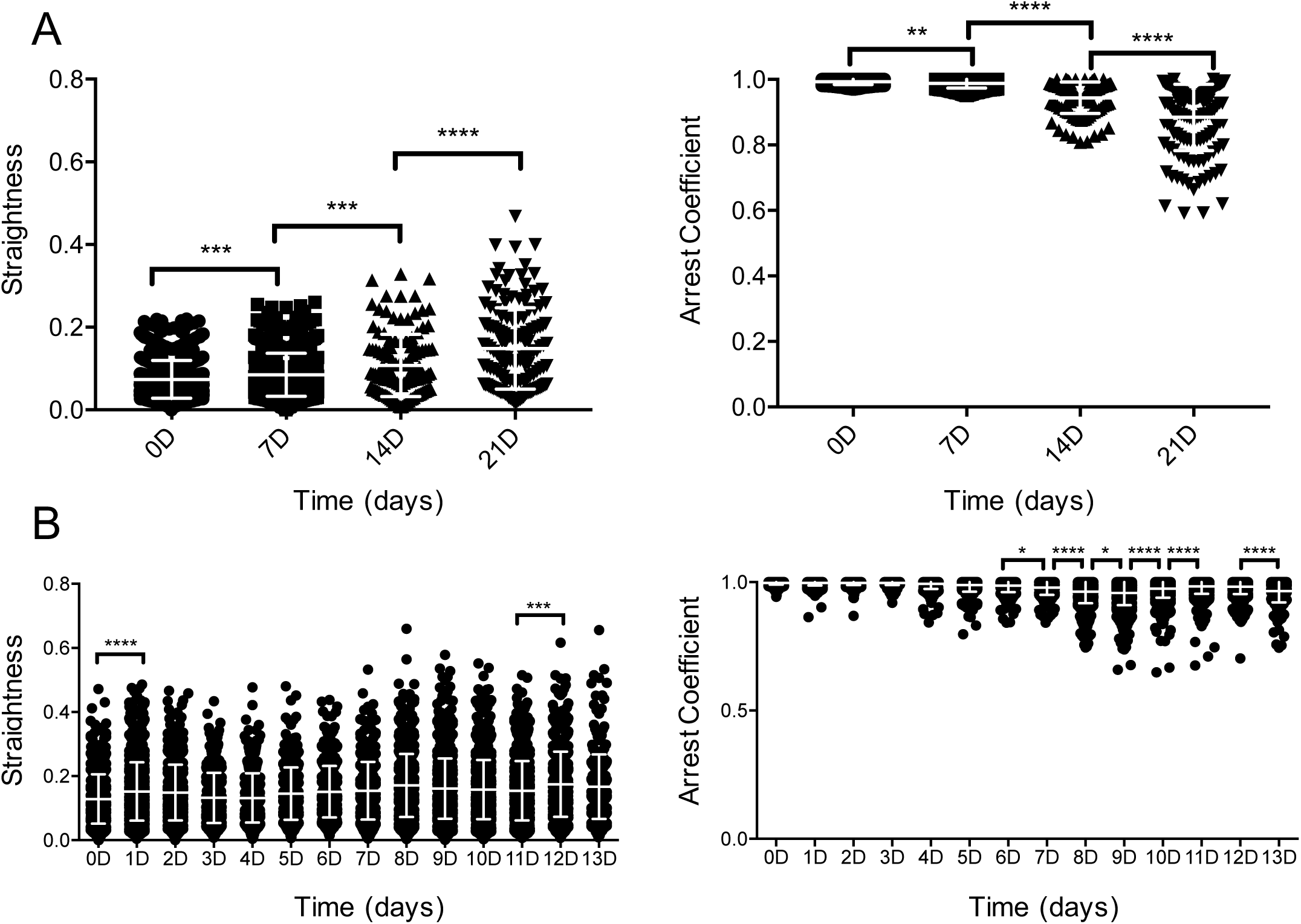
NK cell differentiation leads to increasingly directed migration. A) Straightness and arrest coefficient for NK cell tracks at weekly time points. Error bars indicate s.d. Means with significant differences as analyzed by ordinary one-way ANOVAwith Tukey’s multiple comparison test are shown (**p< 0.01, ***p< 0.001, ****p< 0.0001). n=932 (0D), 803 (7D), 134 (14D), 148 (21D). Straightness and arrest coefficient for NK cell tracks at daily time points. Error bars indicate s.d. Means with significant differences are determined by ordinary one-way ANOVAwith Tukey’s multiple comparison test (*p< 0.05, ***p< 0.001, ****p< 0.0001). Sample size for each individual time point are listed in Materials and Methods. All data shown are representative of 3 independent experiments.

#### Mode of migration depends on NK cell developmental stage

Migrating cells can either exhibit directed motion, constrained motion, or random diffusion (Krummel *et al.*, 2016). Random diffusion represents the type of movement expected of a cell of a certain size due to Brownian motion (Khorshidi *et al.*, 2011). In this case, the mean square displacement (MSD) will be linear with time, whereas directed or constrained tracks will deviate above or below the linear trend, respectively (Krummel *et al.*, 2016). To characterize the diffusivity of our NK cell precursors, we calculated the MSD of NK cell tracks at the weekly time points (Figure 4A). In agreement with our previous measurements of track length and displacement, MSD progressively increased over time, starting at 546.8±549.6 µm^2^ at day 0 for *t*=450 minutes and increasing to 6405.0±10447.7 µm^2^ at day 21 for the same *t* value.

**Figure 4.**
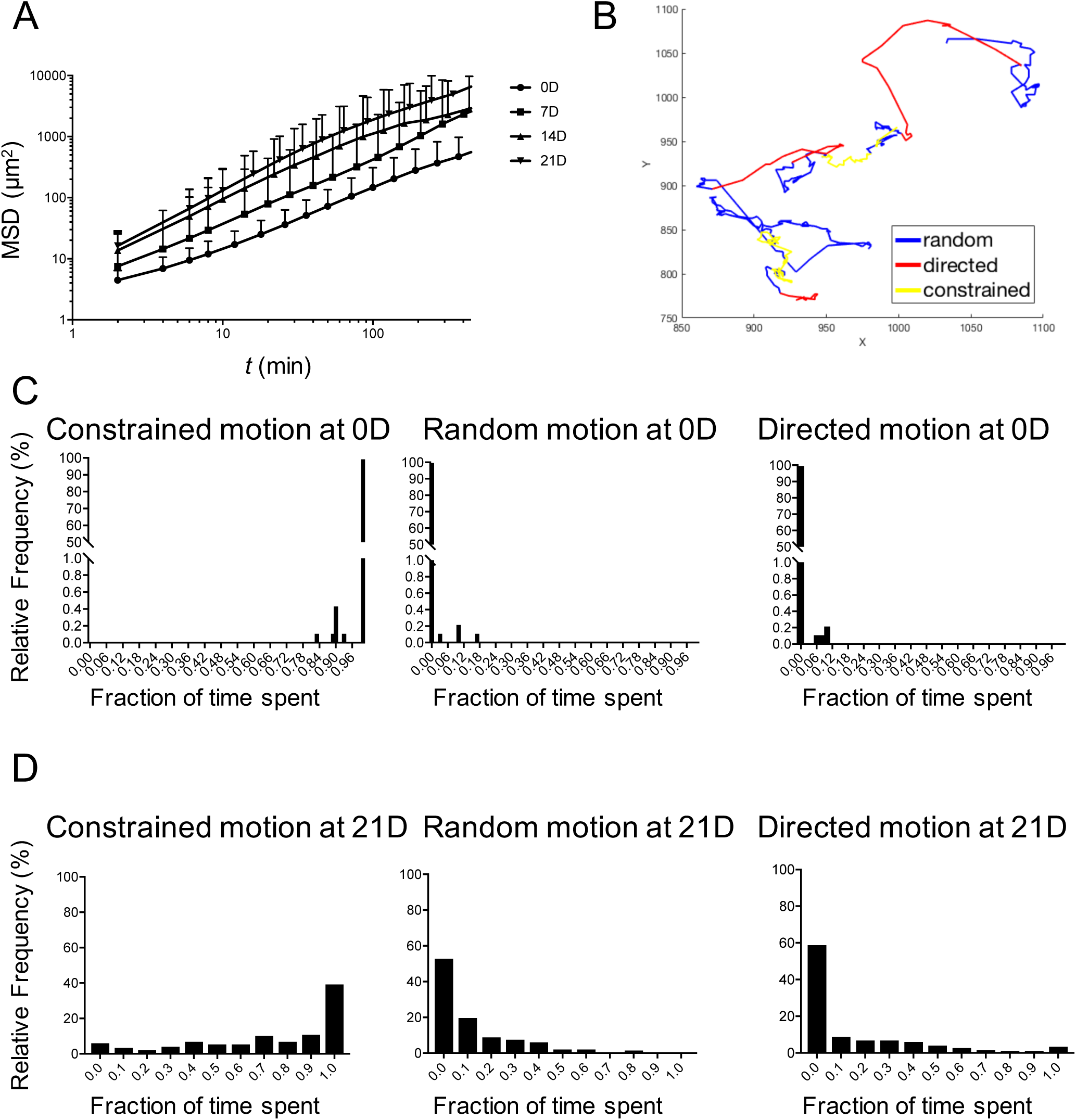
NK cell differentiation is associated with distinct modes of migration. A) Mean square displacement (MSD) of tracks acquired at weekly time points. Graph is truncated at 450 min because few cell tracks persist for longer. Error bars indicate s.d. B) Representative NK cell track after 21 days of development shown with segments corresponding to each migration mode labeled.Fraction of time spent in either constrained, random, or directed motion for each cell after 0 days of development. n=932. D) Fraction of time spent in either constrained, random, or directed motion for each cell after 21 days of development. n=148. All data shown are representative of 3 independent experiments.

While these values reflected a population-based measurement, many cells exhibited complex behaviors with multiple modes contained within a single track. To classify cell tracks by their transient properties, we implemented a previously described method for analyzing NK cell migration (Khorshidi *et al.*, 2011). Using this method for each track we calculated both the diffusion coefficient *D* and the diffusion exponent a, which define the slope and curvature of the MSD curve, respectively. Tracks were classified by mode of migration by thresholding on these values (Figure 4B). Applying this analysis to all tracks for a given time point gave the fraction of time cells spent in either constrained versus directed migration. At day 0, the mean fraction of time cells exhibited constrained motion was 99.9% and the mean fraction of time cells underwent directed motion was a mere 0.04% (Figure 4C). This had changed significantly by day 21, where the mean fraction for constrained motion decreased to 71.9% and the mean fraction for directed migration increased to 15.9% (Figure 4D). These results suggest that the increase in mean speed over developmental stage that was described previously is due to a greater propensity for more mature cells to undergo directed migration. By analyzing the mean squared displacement of cell tracks, we observed significantly more directed walks at later stages of NK cell differentiation. This could reflect a migration strategy adopted by more functional NK cell intermediates to maximize target cell killing.

Many cells measured at later time points exhibited seemingly Lévy-type walks, characterized by periods of extended cell arrest interspersed with short, highly directional movements. CD8^+^ T cells similarly utilize Lévy walks, with computational modeling suggesting that this behavior increases the efficiency of locating target cells compared to a purely random walk (Harris *et al.*, 2012). Inhibiting T cell turning, such as through deletion of the non-muscle myosin motor myosin 1g, leads to decreased detection of rare antigens and supports the idea that an inability to effectively perform Lévy-type walks results in a less efficient search strategy (Gerard *et al.*, 2014). In NK cells, similar transient periods of arrest have been previously described to correspond to the formation of conjugates with target cells and cell-mediated killing, although they can also occur spontaneously (Khorshidi *et al.*, 2011). Interestingly, IL-2-activated NK cells, which have greater cytolytic activity than unstimulated cells, spend much less time in arrest while also forming twice as many cell contact compared to resting NK cells, suggesting that regulation of this behavior correlates with relevant functional differences (Olofsson *et al.*, 2014). Therefore, the acquisition of this specific mode of migration may represent a previously unappreciated component of functional maturation. Alternatively, the seeking behavior that develops may enable the location of key developmental cues that are required for NKDI to proceed through differentiation. It will be of interest to determine whether this behavior is a requirement for, or a product of, NK cell maturation.

#### NK cell maturation is accompanied by increased heterogeneity in NK cell migratory behavior

While we observed a global increase in cell motility over time, we also observed increasing heterogeneity of cell migratory behaviors with progressive NK cell maturation. Plotting the distribution of mean speeds for the 7-day time intervals, we observed that although mean speeds progressively increased over time, later time points still retained a few cells exhibiting very low speeds (Figure 5A). Individual analysis of single cell tracks showed that this increased variation in speed is accompanied by more diversity in migration behaviors (Figure 5A). Increased standard deviation of track velocities at later time points was seen when we measured track statistics from both the weekly and consecutive daily datasets (Figure 5B). Mean standard deviation increased from 0.39±0.11 µm/min at day 0 to 1.07±0.51 µm/min by day 21. This heterogeneity was due in part to the shift towards complex behaviors; at the beginning of tracking, cells exhibited primarily low-speed constrained tracks, whereas at later times there was a greater frequency of behaviors such as tracks resembling Lévy-type super-diffusive walks, characterized by a series of short, constrained motions interspersed with periods of highly directed motion (Harris *et al.*, 2012; Krummel *et al.*, 2016). Additionally, a small subset of mature cells exhibited purely ballistic migration, travelling in essentially a straight line for the duration of migration, particularly at later time points (14 and 21 days).

**Figure 5.**
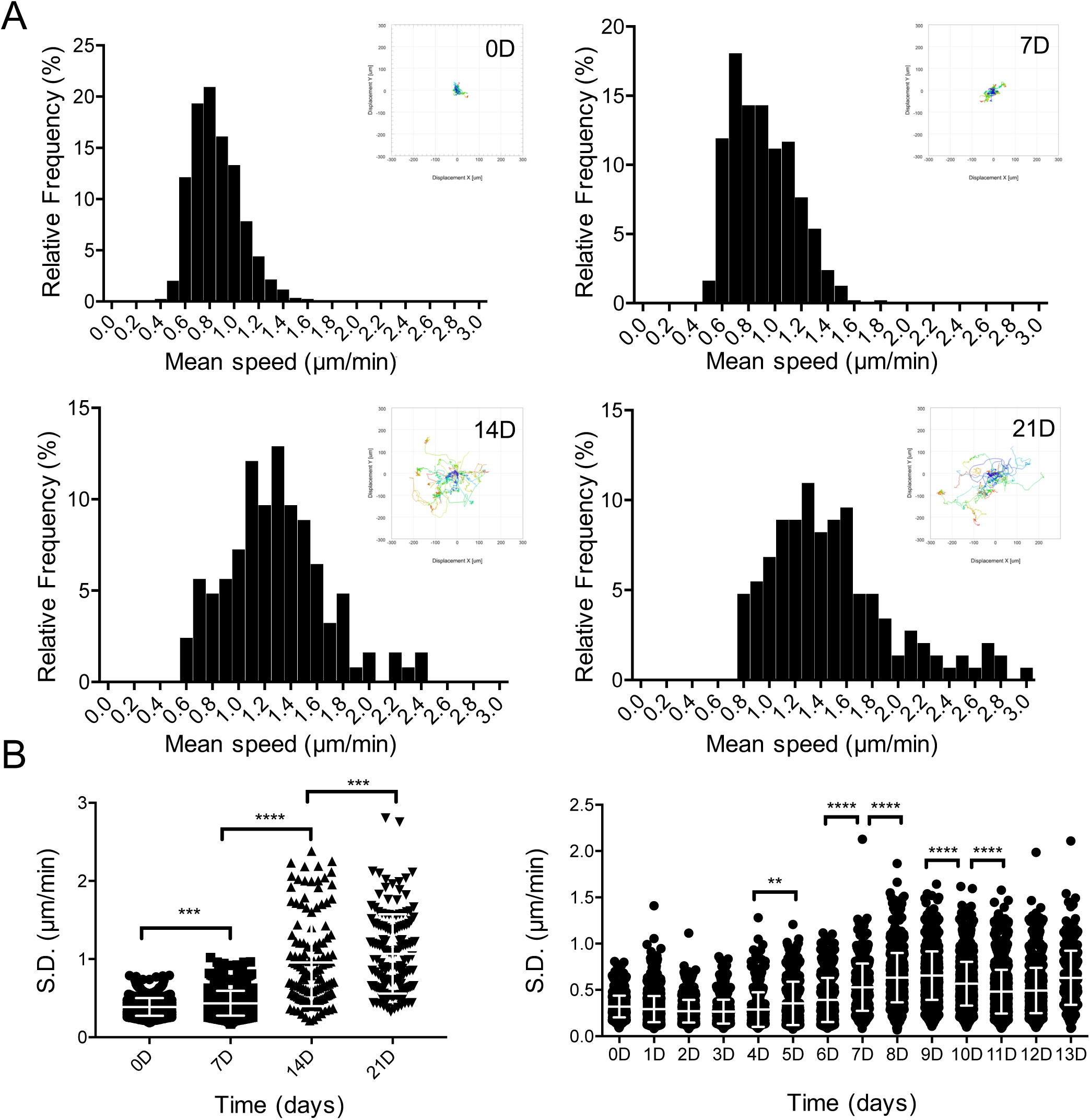
NK cell maturation is correlated with increased heterogeneity of migratory phenotype. A) Histogram of mean speeds observed at the weekly time points. n=932 (0D), 803 (7D), 134 (14D), 148 (21D). Insets: Rose plots of representative tracks. n=20 per graph. B) Standard deviation of the instantaneous speeds observed for each track for daily time points and weekly time points. Error bars indicate s.d. Significance between means determined by ordinary one-way ANOVA with Tukey’s multiple comparisons test (**p< 0.01, ***p< 0.001, ****p< 0.0001). n=932 (0D), 803 (7D), 134 (14D), 148 (21D). All data shown are representative of 3 independent experiments.

### Conclusion

In summary, we have shown that NK cells acquire motility throughout development, progressing from a mostly constrained migratory phenotype to complex walks and more directed migration. The transition to this mode of migration strategy may enable more efficient target cell killing by increasing the rate at which NK cells can conjugate with targets. Additionally, the increased heterogeneity in cell migration at later stages may correspond to NK cells with different functional capabilities, with cells exhibiting greater motility being more effective at cell-mediated cytotoxicity. Continuing to study NK cell dynamics at the single-cell level will likely lead to greater understanding of their development and function.

## Acknowledgments

We thank Dr. Alexandre Carisey for assistance with coding and useful scientific discussion, and Dr. Michael Diehl for critical reading of the manuscript. EL08.1D2 stromal cells were a kind gift from Drs. Jeffrey Miller and Elaine Dzierzak. This work was supported by the Virginia and L.E. Simmons Family Foundation and an American Society of Hematology Junior Faculty Scholar Award to EMM.

## Supplemental movie legends

**Supplementary Movie 1.** Cropped video of a single-cell track 21 days after the initiation of imaging (also shown in Figure 2A). Cells were imaged at 2-minute intervals as described in the text. Scale bar 50 µm.

**Supplementary Movie 2.** Live-cell phase-contrast video of the co-culture 0 days after initiation of tracking. Cells were imaged at 2-minute intervals as described in the text. Scale bar 300 µm.

**Supplementary Movie 3.** Live-cell phase-contrast video of the co-culture 21 days after initiation of tracking. Cells were imaged at 2-minute intervals as described in the text. Scale bar 300 µm.

